# Localized SARS-CoV-2 Infection Triggers a Tissue-Wide Antiviral Response and Functional Impairment of Olfactory Sensory Neurons

**DOI:** 10.64898/2026.09.19.752823

**Authors:** Jiaying Liu, Muhammad Shoaib Akhtar, Hongwei Liu, Yaejin Kim, Brianna M. Ramirez, Anthony L. Weidner, Austin Chan, Maxwell Collins, Stefan Rothenburg, Lark L. Coffey, Qizhi Gong

**Affiliations:** Department of Cell Biology and Human Anatomy, School of Medicine, University of California Davis, United States; Department of Pathology, Microbiology and Immunology, Weill School of Veterinary Medicine, University of California Davis, United States; Department of Microbiology and Immunology, School of Medicine, University of California Davis, United States

**Keywords:** Coronavirus, COVID-19, Cytokine, Interferon-stimulated response, Macrophage, Olfactory epithelium, Olfactory dysfunction

## Abstract

Olfactory dysfunction is a hallmark of COVID-19, yet the mechanisms by which severe acute respiratory syndrome coronavirus 2 (SARS-CoV-2) infection causes widespread sensory impairment remain incompletely understood. We compared the olfactory epithelial tropism of multiple SARS-CoV-2 variants in a mouse model and observed that Alpha and Beta variants exhibited the greatest infectivity for the olfactory epithelium (OE), whereas Omicron rarely infected this tissue despite comparable pulmonary viral titers. We further defined the effects of SARS-CoV-2 on olfactory sensory neurons (OSNs) using immunohistochemistry, single-cell RNA sequencing, spatial gene expression characterization, and functional odor stimulation. The localized infection of sustentacular cells triggered a tissue-wide interferon-stimulated antiviral response that extended beyond infected regions for Alpha and Beta but was largely absent following Omicron infection. Mature OSNs transiently adopted an interferon-responsive state before exhibiting persistent downregulation of odorant signal transduction, mitochondrial, and activity-dependent gene pathways. Consistent with these transcriptional changes, odor stimulation failed to elicit normal activity-dependent gene expression during SARS-CoV-2 infection, indicating impaired neuronal function without widespread neuronal loss. Progressive accumulation of macrophages further indicated sustained inflammatory remodeling of the OE. Together, these findings demonstrate that SARS-CoV-2 infection initiates tissue-wide antiviral signaling in the OE that persistently disrupts OSN function, providing a mechanistic framework for COVID-19-associated anosmia.

## Introduction

The olfactory epithelium (OE) represents a unique neuroimmune interface^1–5^. It is continuously exposed to inhaled pathogens while having to preserve the function of olfactory sensory neurons (OSNs), which are essential for odor perception^3,6–8^. However, the mechanisms by which viral infection disrupts neuronal function within the OE remain poorly understood^9–13^. The high prevalence of olfactory dysfunction observed during the early phase of the COVID-19 pandemic, affecting up to 80% of infected patients, highlighted the clinical significance of the question and provided an unprecedented opportunity to investigate how respiratory viruses alter olfactory sensory function^14–17^.

Severe acute respiratory syndrome coronavirus 2 (SARS-CoV-2), the causative virus of COVID-19, primarily infects sustentacular cells within the OE. Sustentacular cells are non-neuronal cells that provide structural and metabolic support to OSNs. They express the viral entry receptors angiotensin-converting enzyme 2 (ACE2) and transmembrane serine protease 2 (TMPRSS2)^18–23^. In contrast, mature OSNs are rarely infected^9,24,25^. This observation suggests that olfactory dysfunction arises predominantly through indirect mechanisms rather than direct viral infection of sensory neurons. Previous studies have implicated olfactory deciliation, inflammatory responses, alterations in OSN nuclear architecture, and odorant receptor expression in SARS-CoV-2-related anosmia^9,18,19,22,26–28^. While these mechanisms likely contribute to olfactory dysfunction, they do not explain how infection of sustentacular cells is translated into olfactory functional deficits.

Behavioral studies have further demonstrated that SARS-CoV-2-infected animal models exhibit impaired olfactory function^10,19,26,29,30^. However, the prior work does not distinguish whether the olfactory deficits originate within the peripheral OE or result from alterations in downstream olfactory circuits. Moreover, it remains unclear whether antiviral responses are restricted to infected cells or propagate throughout the OE to reprogram neighboring non-infected epithelial and neuronal cells, how these responses evolve over time, and whether they produce persistent neuronal dysfunction.

Here, we combined immunohistochemistry, single-cell RNA sequencing, spatial transcriptomics, and functional odor stimulation in an established murine model to define how SARS-CoV-2 infection remodels the OE. We demonstrate that localized infection initiated a tissue-wide antiviral response extending far beyond infected regions, induced a transient interferon-responsive state in mature OSNs, and was followed by persistent suppression of odor transduction and activity-dependent gene programs. These transcriptional changes were accompanied by impaired odor-evoked neuronal activation and progressive inflammatory remodeling despite preservation of neuronal abundance. Together, our findings identify tissue-wide bystander antiviral signaling as a mechanism linking SARS-CoV-2 infection to persistent OSN dysfunction.

## Results

### Variant-dependent differences in SARS-CoV-2 pathogenesis and tropism in the OE

The K18-hACE2 transgenic mouse model enables SARS-CoV-2 infection and is used to study disease pathogenesis^10,31–33^. The evolution of SARS-CoV-2 and its differential impact on olfactory function make it essential to understand viral interactions with the olfactory mucosa. To understand how different SARS-CoV-2 variants infect olfactory mucosa, we intranasally inoculated K18-hACE2 mice with B.1 (WA1-like), Alpha (B.1.1.7), Beta (B.1.351), Delta (B.1.617.2), Omicron (B.1.1.529), or phosphate buffered saline (PBS) mock control (Figure 1a). Mice inoculated with all variants except Omicron lost weight starting at 2 dpi. Animals infected with B.1 and Delta variants lost 20% or more of their starting weight by 6 dpi, which met euthanasia criteria (Figure 1b).

**Figure 1.**
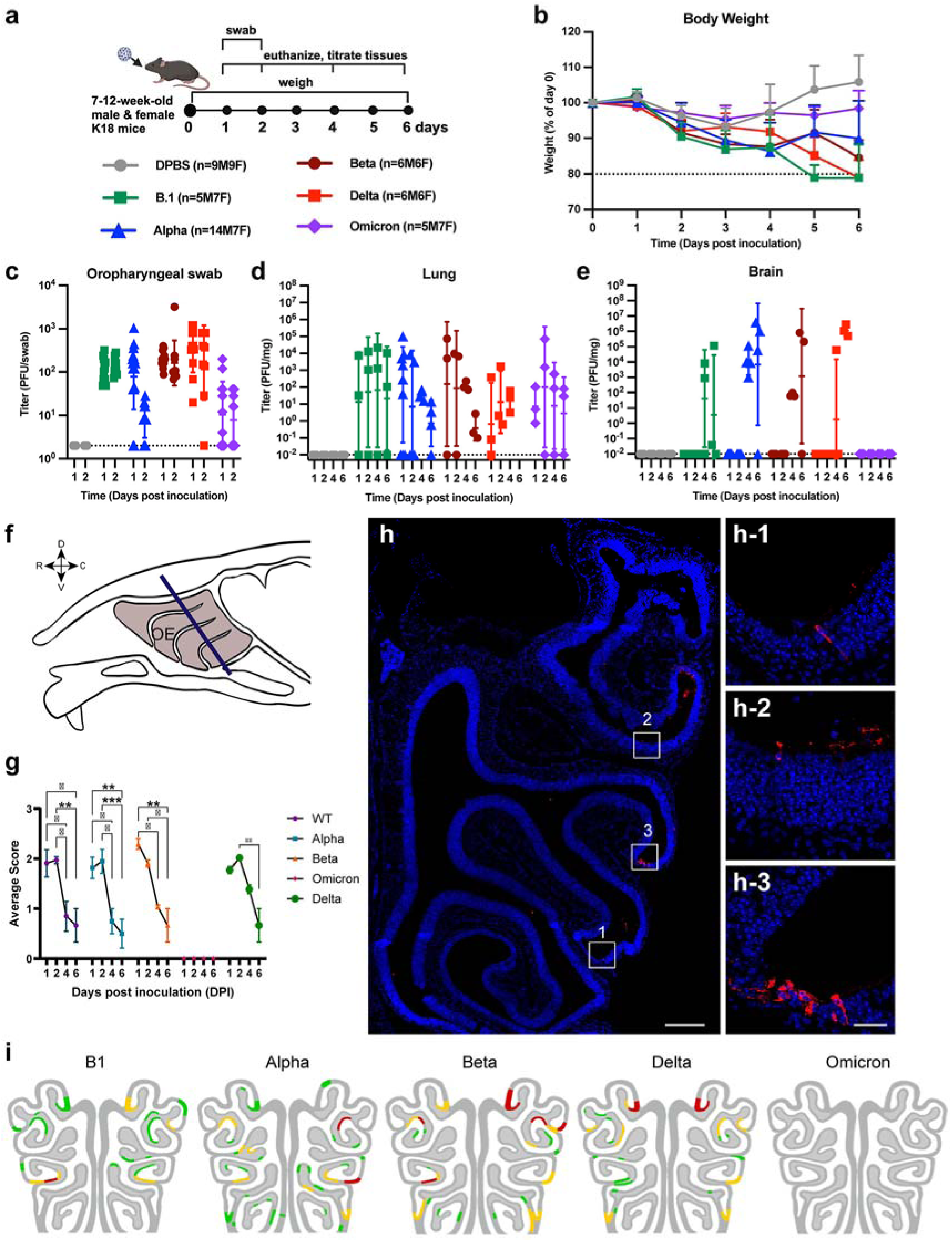
SARS-CoV-2 variants of concern show distinct pathogenesis, tissue tropism and olfactory epithelium infection patterns in K18-hACE2 mice. **a** Study design. Seven- to 12-week-old male and female K18-hACE2 mice were inoculated intranasally with 10⁴ or 10^5^ PFU of SARS-CoV-2 lineage B.1, Alpha, Beta, Delta or Omicron, or mock-inoculated with DPBS. Body weight was monitored daily, oropharyngeal swabs were collected on days 1 and 2 post-inoculation (dpi), and subsets of mice were euthanized on 1, 2, 4 and 6 dpi for tissue collection. Group sizes consist of male (M) and female (F) mice. **b** Body weight change over the course of infection, expressed as a percentage of weight on day 0. **c** Infectious virus titers in oropharyngeal swabs collected on 1 and 2 dpi. **d** Infectious virus titers in lung homogenates on 1, 2, 4 and 6 dpi. **e** Infectious virus titers in brain homogenates on 1, 2, 4 and 6 dpi. For b–e, data are presented as mean ± SD; dotted lines in c–e indicate the limit of detection. Group sizes are as in **a**. **f** Schematic sagittal section of the mouse head indicating the position of the olfactory epithelium (OE, shaded) and the coronal plane (blue line) used for sectioning in **g**–**i**. D, dorsal; V, ventral; R, rostral; C, caudal. **g** Line graph of average OE infection scores of variants at 1 dpi, 2 dpi, 4 dpi, and 6 dpi (n = 3 per variant per time point). NP score (as in h) for OEs with infection sites: h-1 = 1, h-2 = 2, h-3 = 3; and for OEs without infection sites is 0. Data is Mean ± SEM \**p* < 0.05, \*\**p*< 0.01, **** *p* < 0.0001 (ANOVA Mixed Effects Analysis). **h** Representative immunofluorescence images of nasal turbinates (1 dpi; NP, red; DAPI, blue) illustrating the scoring system for NP signal: 1–2 infected cells (h-1); 3–5 cells (h-2); >5 cells (h-3); no infection, score 0. Scale bars, 200 µm (h) and 30 µm (h1-3). **i** Heatmaps of nasal epithelium infection patterns per SARS-CoV-2 variant (1dpi; n = 3 per variant), showing average NP score (as in h) per region of interest, color-coded: green, as in h-1; yellow, as in h-2; red, as in h-3.

All variants were detected in oropharyngeal swabs in most mice 1 and 2 dpi and in lung 1, 2, 4 and 6 dpi with a general decreasing trend in lung viral titers starting at 2 dpi (Figure 1c-d). We further investigated SARS-CoV-2 infection in the brain due to the reported hACE2 expression in K18-hACE2 mice^34,35^. Brain titers were undetectable at 1-2 dpi for all variants. Alpha, Beta, and Delta variants showed increasing brain titers by 4-6 dpi. B.1 showed modest and stable titers; Omicron showed no brain infection throughout the time course (Figure 1e). These results align with prior findings on Omicron’s minimal neurotropism^31,36^.

To evaluate the spatial and temporal pattern of SARS-CoV-2 infection within the K18-hACE2 mouse OE, we collected coronal sections from animals at 1–6 dpi (Figure 1f-g). Human ACE2 expression in the OE was in the sustentacular cells located on the apical surface (supplemental Figure 1). The viral nucleocapsid protein (NP) was detected by immunostaining to identify infected cells. Consistent with published findings^10,18,20^, viral NP was detected primarily in sustentacular cells. NP signals were localized primarily within nasal turbinates and were not detected in the septum across all variants, with no clear strain-specific spatial tropism. Except for Omicron, which showed minimal NP signal, infection sites of other variants were sparsely distributed either as a single NP-positive cell or small clusters (Figure 1h). B.1, Alpha, and Beta infection at 10^4^ plaque forming unit (PFU) and Delta infection at 10^5^ PFU show similar levels of infectivity of the OE across time from 1 dpi to 6 dpi (Figure 1g). Beta infection peaked at 1 dpi whereas B.1, Alpha, and Delta peak at 2 dpi and decline afterwards (Figure 1g). Omicron exposure resulted in no detectable infection at any point. For the other variants, multicell infection clusters have largely resolved by 4 dpi and completely resolved by 6 dpi (supplemental Figure 2). Accordingly, OE infection declined significantly at these time points, as measured using a semiquantitative infection score (1, a single infected cell; 2, two to three infected cells; and 3, four or more infected cells (Figure 1g).

### Rapid transcriptional responses in SARS-CoV-2-infected olfactory mucosa

To identify tissue level transcriptomic changes under SARS-CoV-2 infection, we next performed bulk RNA sequencing on olfactory mucosa collected from mice inoculated with Alpha variant, one of the infectious viruses in the OE, or PBS mock control at 1 and 2 dpi (n = 3 biological replicates per group; Figure 2a, supplemental Figure 3). SARS-CoV-2 Alpha variant infection elicited a rapid and robust transcriptional response. A total of 513 genes were differentially expressed (adj. *p* < 0.05) at 1 dpi, including 493 upregulated and 20 downregulated transcripts (Figure 2a). By 2 dpi, the transcriptional response had decreased, with 164 genes remaining differentially expressed, including 144 upregulated and 20 downregulated genes (supplemental Figure 3).

**Figure 2.**
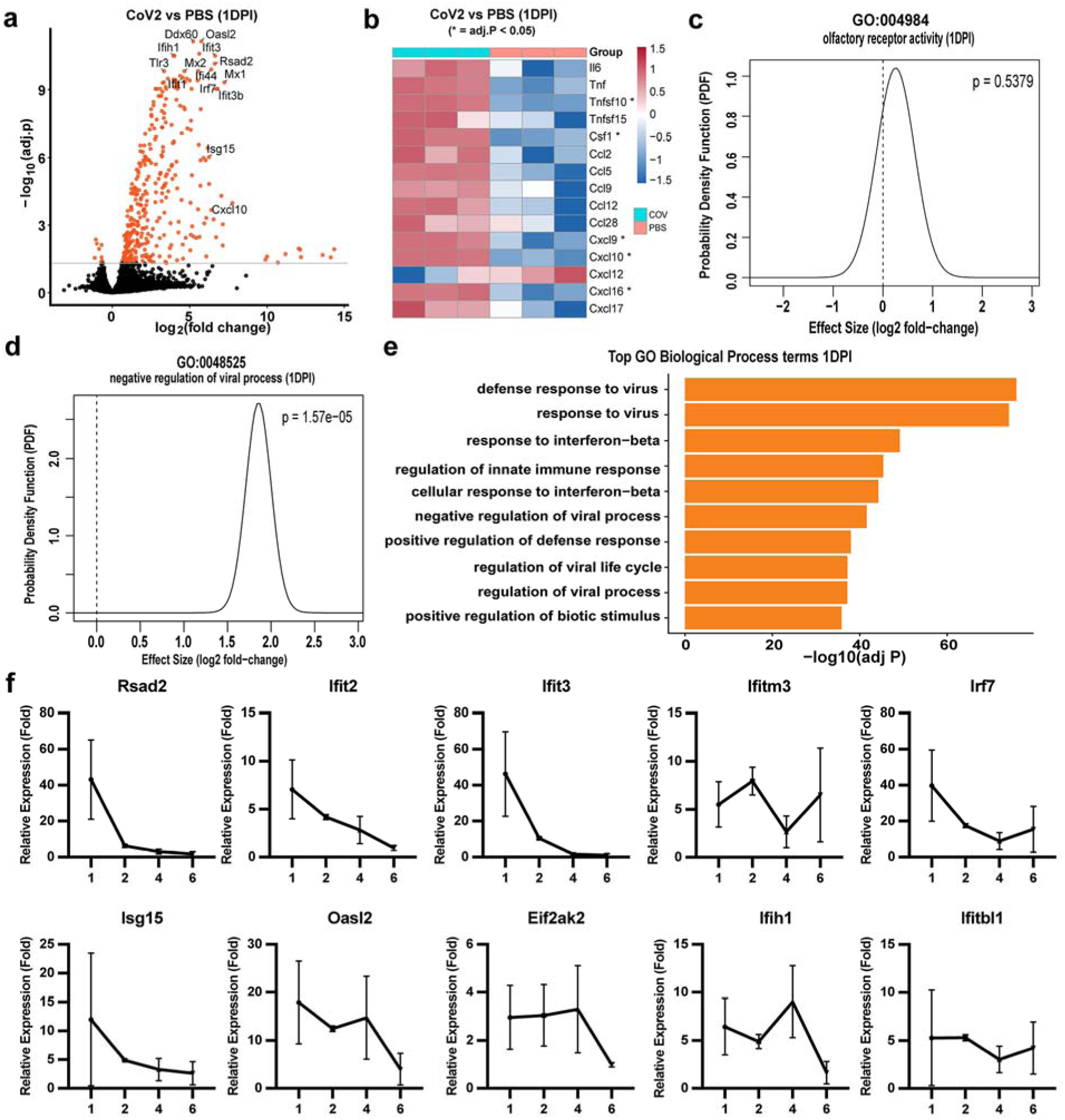
SARS-CoV-2 infection induces a robust antiviral transcriptional response in the olfactory mucosa of K18-hACE2 mice. **a** Volcano plot of differentially expressed genes in bulk olfactory mucosa from SARS-CoV-2 (Alpha)-infected versus PBS mock-infected K18-hACE2 mice at 1 dpi (n = 3 per group). Red, genes with adj. *p* < 0.05 (grey line); selected interferon-stimulated genes are labeled. **b** Heatmap of relative expression (z-score) of cytokine and chemokine transcripts in SARS-CoV-2 versus PBS mock-infected mice (n = 3 per group), from the same dataset as in **a**. Asterisks mark genes significant at 1 dpi (adj. *p* < 0.05). **c**, **d** QuSAGE gene-set analysis of GO:0004984, olfactory receptor activity (**c**) and GO:0048525, negative regulation of viral process (**d**) gene sets. Curves show the probability density function (PDF) of the gene-set effect size (log2 fold-change) relative to the null (dashed line); the olfactory signaling gene set showed no significant shift (pdf > 0.5), whereas the antiviral response gene set showed a significant upward shift (*p* = 1.57 x 10^-5^). **e** Top 10 enriched Gene Ontology biological process (GO BP) terms among genes upregulated in SARS-CoV-2 versus PBS-infected mice, ranked by −log10(adj. *p*). **f** qPCR validation of interferon-stimulated genes *(Rsad2, Ifit2, Ifit3, Ifitm3, Irf7, Isg15, Oasl2, PKR, Ifih1, Ifnb1*) in olfactory mucosa at 1, 2, 4 and 6 dpi, expressed as fold change relative to PBS mock-infected mice. Data are Mean ± SD.

Among the earliest induced transcripts were cytokines, chemokines, and interferon-stimulated genes associated with antiviral immunity. Pro-inflammatory cytokines, including *Il6*, *Tnf*, *Csf1*, *Tnfsf10*, and *Tnfsf15*, together with chemokines from both the *CCL* (*Ccl2*, *Ccl5*, *Ccl9*, *Ccl12*, *Ccl28*) and *CXCL* (*Cxcl9*, *Cxcl10*, *Cxcl12*, *Cxcl16*, *Cxcl17*) families, were evaluated at 1 dpi. Among those, *Tnfsf10, Cxcl10, Cxcl16, Cxcl9*, and *Csf1* showed significant upregulation (Figure 2b, adj. *p* < 0.005). By 2 dpi, expression of most inflammatory mediators had returned toward baseline, with significant upregulation persisting only for *Cxcl10* and *Tnfsf10*.

Previous studies have reported conflicting findings regarding SARS-CoV-2-induced downregulation of olfactory receptor transcripts^9,19,37^. Because olfactory receptor genes exhibit substantial inter-individual variability, we evaluated their expression using both gene-level differential expression analysis (Limma-Voom) and gene set enrichment analysis (QuSAGE)^38^. No olfactory receptor genes were significantly downregulated (adj. *p* < 0.05; log2FC < -1), whereas only nine of the more than 1,000 olfactory receptor genes were significantly upregulated. Likewise, gene set analysis of the olfactory receptor activity pathway (GO:0004984) detected no significant changes at either 1 or 2 dpi (Figure 2c, supplemental Figure 3b). These findings indicate that early olfactory and behavioral deficits in the K18-hACE2 model were unlikely to result from global suppression of olfactory receptor gene expression.

Functional enrichment analysis demonstrated that the early transcriptional response was dominated by antiviral signaling. Gene Ontology analysis identified 187 significantly enriched biological process terms among upregulated genes at 1 dpi (*adj. p < 0.05*; supplemental table 1), with ‘defense response to virus’ representing the most significantly enriched category (Figure 2e). Consistent with this observation, QuSAGE analysis confirmed robust enrichment of the antiviral gene set (GO:0048529) at 1 dpi (Figure 2d). Although pathway-level enrichment was reduced by 2 dpi (Supplemental Figure 3c, supplemental table 2), many individual antiviral genes remained elevated (Figure 2f), indicating rapid activation followed by partial resolution of the antiviral transcriptional program.

To further define the kinetics of this response, representative antiviral genes were quantified by RT-qPCR at 1, 2, 4, and 6 dpi (n = 3 biological replicates per group). *Rsad2, Ifit3, Irf7, Isg15,* and *Ifit2* exhibited the strongest induction at 1 dpi and declined progressively thereafter. In contrast, *Oasl2, Eif2ak2* (encoding PKR)*, Ifih1, Ifit1bl1,* and *Ifitm3* showed more sustained expression, with several remaining elevated through 4 dpi before returning toward baseline by 6 dpi (Figure 2f). Together, these findings demonstrate that SARS-CoV-2 infection induced a rapid antiviral transcriptional response in the olfactory mucosa that was downregulated at the bulk tissue level within the first week after infection.

### Localized infection elicits tissue-wide antiviral responses in the OE

To investigate whether antiviral responses were restricted to sites of SARS-CoV-2 infection, we performed multiplex RNAscope in situ hybridization to simultaneously detect viral RNA and interferon-stimulated gene (ISG) transcripts within the OE. Consistent with the distribution of NP immunostaining shown earlier (Figure 1), SARS-CoV-2 spike transcripts were confined to discrete foci (Figure 3a). Six ISGs were examined, selected from top differentially expressed ISG transcripts identified in the bulk RNA-seq including *Rsad2, Isg15, Ifit3, Ifih1, Irf7,* and *Eif2ak2* (Figure 3b-g). Compared with the minimal expression observed in mock-infected controls, all six ISGs were robustly induced throughout the OE. ISG expression was detected in both sustentacular cells and OSNs, extending beyond sites of detectable viral transcript.

**Figure 3.**
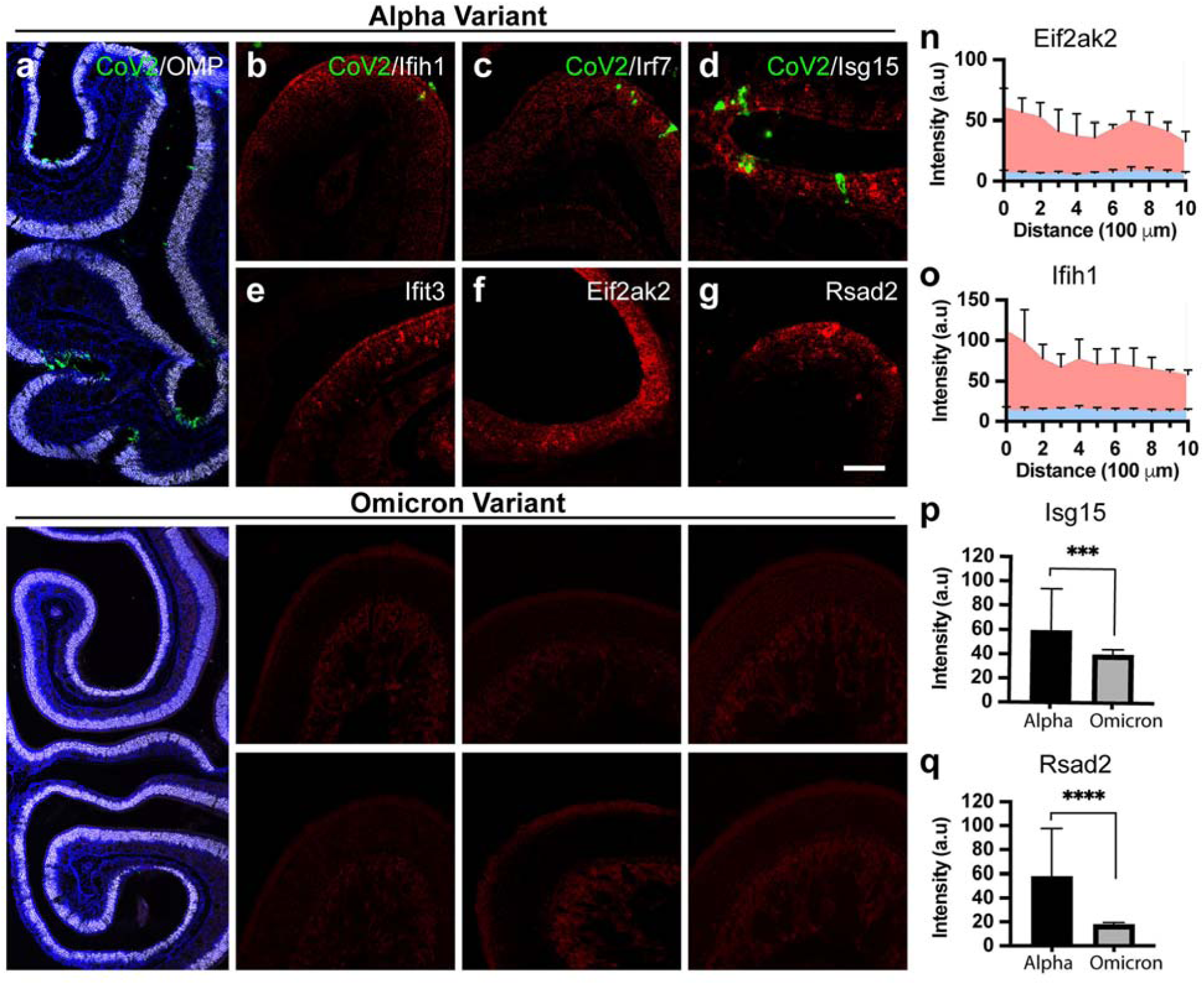
Local SARS-CoV-2 infection induces tissue-wide interferon-stimulated gene response. **a, b** Low-magnification RNAscope images of the OE co-stained for *SARS-CoV-2* (CoV2, green) and *Omp* (white) mRNA, with nuclei (DAPI, blue), from Alpha- (**a**) and Omicron-infected (**b**) K18-hACE2 mice at 1 dpi, showing abundant viral RNA in Alpha-but minimal viral RNA in Omicron-infected tissue. **b–g** RNAscope images of interferon-stimulated genes *Eif2ak2* (**b**), *Rsad2* (**c**), *Isg15* (**d**), next to CoV2 signal (green), *Ifit3* (**e**), *Ifih1* (**f**) and *Irf7* (**g**) (red), away from CoV2 signal, in the OE of Alpha-infected mice**. h–m** As in **b–g**, for Omicron-infected mice. **n, o** Quantification of Eif2ak2 (**n**) and Ifih1 (**o**) RNAscope signal intensity as a function of distance from the infection focus in Alpha-infected (red) versus PBS mock-infected (blue) mice, in 100 µm bins out to 1 mm. Signal in infected tissue approaches the PBS baseline by ∼1 mm from the infection site. Data are mean ± SD. **p, q** Quantification of *Isg15* (**p**) and *Rsad2* (**q**) RNAscope signal intensity within the infected region of interest in Alpha (A)-versus Omicron (O)-infected mice. \*\*\**p* < 0.001, \*\*\*\**p* < 0.0001 (unpaired t-test).

At 1 dpi, when antiviral responses were readily detected according to bulk RNA-seq and RT-qPCR analysis, viral RNA remained confined to localized infection foci, whereas ISG transcripts were distributed broadly throughout the OE. Notably, ISG expression was strongest in cells immediately adjacent to infection foci, and gradually decreased with increasing distance, forming a spatial expression gradient (Figure 3n,o). Regions infected by SARS-CoV-2 displayed reduced *Omp* transcript abundance, consistent with the previous report^9^ (Figure 3a).

To determine whether this widespread antiviral response resulted from local OE infection or systemic SARS-CoV-2 infection, we examined mice infected with the Omicron variant. We did not detect Omicron infection in the OE, despite its pulmonary infectivity being comparable to that of the Alpha variant (Figure 1d). In contrast to Alpha infection, expressions of *Eif2ak2*, *Rsad2*, *Isg15*, *Ifit3*, *Ifih1*, and *Irf7* remained minimal in the OE of Omicron-infected animals and consistent with the levels observed in mock-infected OE (Figure 3h-m, supplemental Figure 6). Quantitative analysis demonstrated significantly greater antiviral gene expression in Alpha-infected than Omicron-infected OE (Figure 3p-q, *p* < 0.001). Together, these findings indicate that robust antiviral activation within the OE was closely associated with productive local infection rather than systemic SARS-CoV-2 infection alone. The widespread distribution of ISG transcripts further suggests that antiviral responses were not restricted to infected cells, prompting us to investigate cell-type-specific responses by single-cell RNA sequencing (scRNA-seq, see below).

### SARS-CoV-2 induces a transient ISG-high neuronal state and persistent transcriptional remodeling

To investigate cell-type-specific responses in the olfactory mucosa, we performed scRNA-seq on SARS-CoV-2-Alpha-variant inoculated and mock-inoculated animals (n = 3 per group) at 1 and 6 dpi. Following quality control, 110,874 and 119,308 cells were retained for analysis at 1 and 6 dpi, respectively (Figure 4a). We identified major cell types of the olfactory mucosa: mature OSNs (mOSNs), immature OSNs (iOSNs), immediate neuronal precursors (INPs), globose basal cells (GBCs), horizontal basal cells (HBCs), sustentacular cells (SUSs), Bowman’s gland cells (BGs), respiratory epithelial cells (REs), fibroblasts (FCs), olfactory ensheathing cells (OECs), immune cells (IMCs), and endothelial cells (ECs) (Figure 4b, supplemental Figure 4).

**Figure 4.**
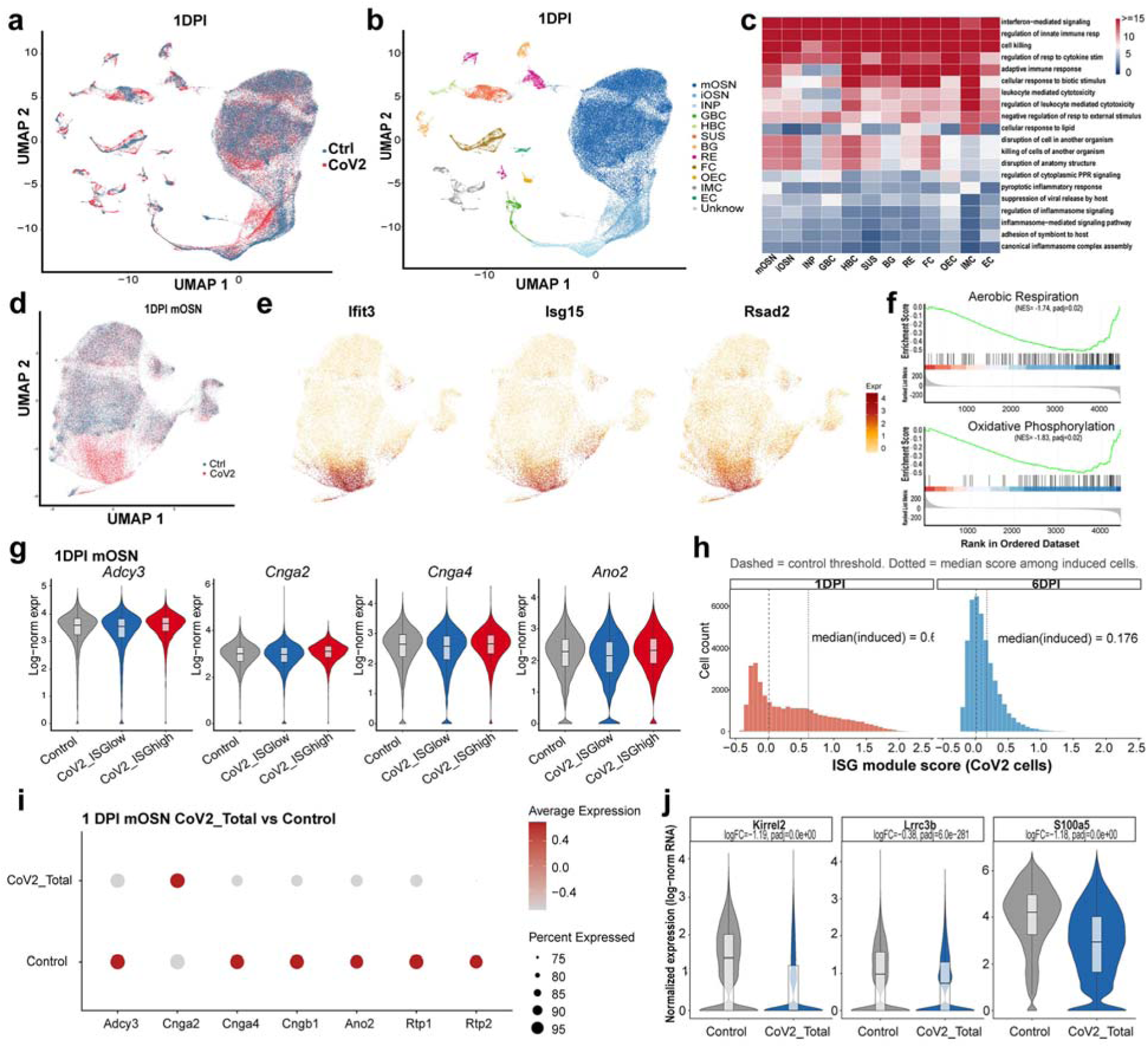
Single-cell RNA-seq reveals interferon-stimulated gene induction and suppression of olfactory transduction genes in mature OSNs. **a** UMAP of olfactory mucosa scRNA-seq from Alpha-infected and control K18-hACE2 mice at 1 dpi, colored by cell type: mature olfactory sensory neuron (mOSN), immature OSN (iOSN), immediate neuronal precursor (INP), globose basal cell (GBC), horizontal basal cell (HBC), sustentacular cell (SUS), Bowman’s gland (BG), respiratory epithelium (RE), fibroblast (FC), olfactory ensheathing cell (OEC), immune cell (IMC) and endothelial cell (EC). **b** UMAP as in **a**, colored by condition (control, CoV2). **c** Heatmap of GO biological process terms upregulated in CoV2-versus control of 1 dpi across all cell types (adj.*p* < 0.05, log2FC > 0.5), simplified per cell type (Wang similarity 0.7) and colored by −log10(adj.*p*). **d** UMAP of mOSN cells only, colored by condition. **e** Expression of *Ifit3, Isg15* and *Rsad2* in mOSN cells from CoV2-infected mice only. **f** GSEA of GO BP terms (aerobic respiration, oxidative phosphorylation) comparing ISGhigh versus control mOSN cells, with the top 24 interferon-stimulated genes (ISGs) defining the ISG module score excluded from the ranked list (stripTop24) to test effects independent of the core ISG signature. NES and adjusted P values are indicated. **g** Expression of olfactory signal-transduction genes (*Adcy3, Cnga2, Cnga4, Ano2*) in mOSN cells from control mice and CoV2-infected mice split into ISGlow and ISGhigh classes at 1 dpi. **h** Distribution of the top24 ISG module score in CoV2-infected cells at 1 and 6 dpi. Dashed line, induction threshold derived from control cells; dotted line, median score among induced cells. Percentage of induced cells and median score among induced cells are indicated. **i** Dot plot of core olfactory transduction-cascade genes in CoV2-infected (total) versus control mOSNs at 6 dpi. Dot color, scaled average expression; dot size, percentage of cells expressing the gene. **j** Expression of mOSN activitity dependent genes (Kirrel2, Lrrc3b, S100a5) in control versus CoV2-infected (total) mOSN cells at 6 dpi. log2 fold-change and adjusted P values (Wilcoxon rank-sum test) are indicated.

Differential gene expression analysis was performed for each cell type to determine whether specific cell populations in the olfactory mucosa respond to SARS-CoV-2 infection. Gene Ontology Biological Process (GO BP) enrichment of upregulated differentially expressed genes (BH-adj. *p* < 0.05) identified the interferon-mediated signaling pathway (GO:0140888) as the most significantly enriched pathway across all cell types at 1 dpi (adj. *p* < 1 x 10^-14^), along with additional GO BP terms associated with innate and adaptive antiviral signaling (Figure 4c). These findings indicate that the acute antiviral transcriptional program was broadly induced throughout the olfactory mucosa, rather than being confined to IMCs or sustentacular cells, which were directly infected by SARS-CoV-2. The magnitude of SARS-CoV-2-induced response was highly cell-type dependent. The fraction of significantly differentially expressed genes ranged from 1.3% in GBCs, to 2.8% in mOSNs, and to 32.2% in IMCs, the highest of any cell type at 1 dpi (3606/11,203 genes tested, supplemental Figure 4d). Pathway breadth followed the same pattern with IMCs showing the greatest number of enriched GO BP terms (Figure 4c and supplemental Figure 4c).

Cells from SARS-CoV-2-inoculated and mock control samples showed largely similar distributions in the overall UMAP embedding (Figure 4b). Further clustering of the 1 dpi mOSNs revealed a distinct group present predominantly in the infected animals (Figure 4d). This cluster was characterized by the upregulation of ISGs, including *Ifit3*, *Isg15*, and *Rsad2* (Figure 4e, S4). This subpopulation accounted for 29.7% of all mOSNs in infected animals at 1 dpi, whereas the ISG-high population was nearly absent (0.05%) in mock-infected controls. Differential expression of ISGs was also detected in the ISG-low mOSNs in the infected samples (Figure 4h).

We subsequently compared transcriptional profiles between ISG-high and ISG-low cells relative to controls. Beyond interferon signatures, extensive transcriptional profile shifts were observed in mOSNs. ISG-high mOSNs showed coordinated suppression of oxidative metabolism. After exclusion of the ISG gene panel (see Methods), gene set enrichment analysis (GSEA) identified oxidative phosphorylation (GO:0006119; NES = -1.80, adj. *p* < 0.05) and aerobic respiration (GO:0009060; NES = -1.74, adj. *p* < 0.05) among the most significant of the 10 significant GO BP terms (Figure 4f). Olfactory signal transduction genes (*Adcy3*, *Cnga2*, *Cnga4* and *Ano2*) showed biologically insignificant alterations (log2FC < ± 0.25) in both ISG tiers at 1 dpi.

To determine whether the 1 dpi signature diminished over time, we jointly scored the same ISG modules across independently clustered and tiered 1 dpi and 6 dpi mOSN datasets. The breadth of induction, measured by ISG module scores, showed a similar percentage of induced cells, but the amplitude among induced cells collapsed, with the median module score falling from 0.614 to 0.176 (Figure 4h). At the functional level, 6 dpi mOSNs continued to show suppression of oxidative phosphorylation (GO:0006119; NES = -1.70, adj. *p* < 0.05), replicating the 1 dpi finding. In addition, two unique programs were identified in 6 dpi mOSNs, cilium loss and proteostatic stress (supplemental Figure 4). Some olfactory signal transduction genes showed significant downregulation, including *Cnga4* (logFC = -0.88), *Cngb* (logFC = - 0.84), *Rtp1* (logFC = - 0.49), and *Rtp2* (logFC = - 1.12) (Figure 4i). In addition, olfactory activity-dependent genes, *S100a5*, *Kirrel2,* and *Lrrc3b*, though not significantly changed at 1 dpi, were all significantly downregulated in mOSNs from SARS-CoV-2-infected animals at 6 dpi^39,40^ (Figure 4j). Together, these findings reveal a biphasic mOSN response to SARS-CoV-2, characterized by an acute ISG-high state with suppressed oxidative metabolism followed by attenuation of interferon signaling but persistent metabolic dysfunction and emerging defects in olfactory activity–dependent programs.

### Infection of the OE correlated with transient and extensive macrophage accumulation

Macrophages are key effectors of the innate immune response and have been proposed to support both neuronal maintenance and pathogen clearance, either of which could influence olfactory function in the OE^2,41,42^. To determine whether SARS-CoV-2 infection elicits macrophage recruitment, we quantified macrophage infiltration by Iba1 immunostaining in the OE of SARS-CoV-2 Alpha-inoculated mice (Figure 5a). The quantitative analysis was done by sampling three regions: the virally infected turbinate, an adjacent non-infected turbinate, and the nasal septum, which is rarely infected (Figure 5a). Comparing macrophage responses across these regions allowed us to distinguish a localized response from one that extends throughout the olfactory mucosa.

**Figure 5.**
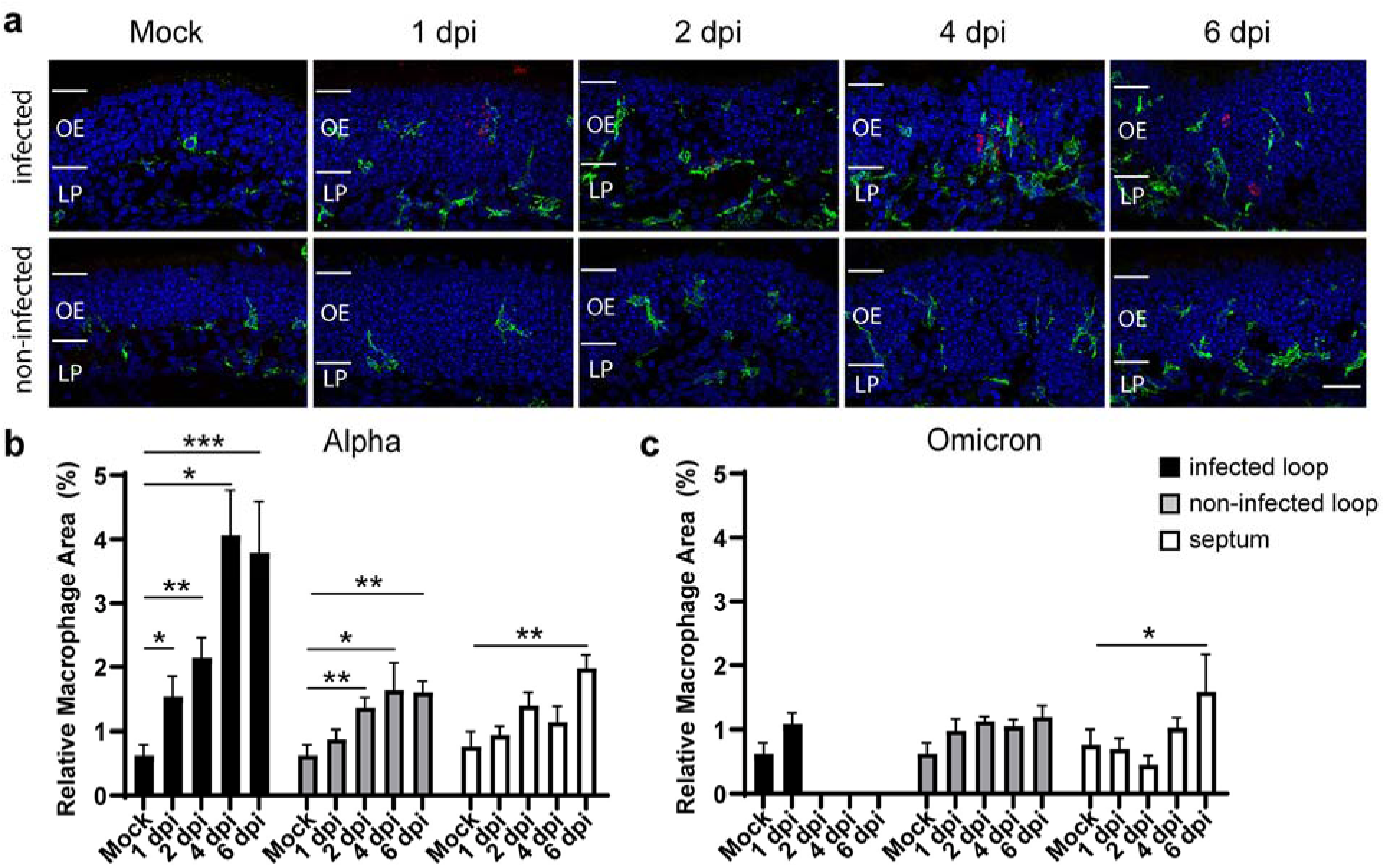
Local macrophage infiltration dynamics in the olfactory mucosa. **a** Representative immunofluorescence images of Iba1+ macrophages (green), nucleocapsid protein (NP). Mock OE regions were selected from turbinates at comparable locations. Infected areas, identified by the presence of NP (red), and non-infected turbinate, which does not contain NP signal, are from Alpha-infected mice at 1, 2, 4 and 6 dpi. OE, olfactory epithelium; LP, lamina propria. Scale bars, 30 µm. **b, c** Quantification of relative Iba1+ macrophage area (%) in infected turbinate loops, non-infected turbinate loops, and the septum (rarely infected) over time in Alpha-(b) and Omicron-infected (c) mice. Infected and non-infected loops are from the same CoV2-infected mice; Mock, PBS inoculated mice. \**p* < 0.05, \*\**p* < 0.01, \*\*\**p* < 0.001 versus mock within the comparable region (unpaired t-test).

In mock-inoculated mice, Iba1-positive macrophages were sparse and largely restricted to the basal lamina of the OE, with a ramified morphology (Figure 5a). Following infection, macrophage abundance and process ramification increased initially only within the infected sites at 1 dpi (*p* < 0.05). By 2 dpi, macrophage infiltration became also evident in non-infected turbinate, and the Iba1-positive area increased approximately threefold in the infected turbinate (0.63 ± 0.16% vs. 2.15 ± 0.31%, *p* < 0.01) and approximately twofold in the non-infected turbinate (0.63 ± 0.16% vs 1.37 ± 0.15%, *p* < 0.01) relative to controls. By 4 dpi, macrophage occupancy continued to increase in the infected turbinate, whereas the non-infected turbinate maintained a level of infiltration comparable to that observed at 2 dpi. Elevated macrophage abundance persisted through 6 dpi. The nasal septum is the region most distant from the primary site of infection. There was also a significant increase of approximately twofold in Iba1-positive area in the nasal septum (0.77 ± 0.24% vs. 1.98 ± 0.21%, *p* < 0.01) (Figure 5b). Together, these findings indicate that macrophage recruitment was initiated at sites of viral infection and subsequently extended into neighboring uninfected regions of the OE.

In contrast, Omicron infection elicited a markedly attenuated macrophage response. Because Omicron infection produced no detectable infection in the OE beyond 1 dpi, infected turbinates could not be reliably sampled at later time points. Macrophage abundance in non-infected turbinates remained near baseline throughout the time course, whereas septal macrophage area increased only modestly, reaching statistical significance related to controls only at 6 dpi (*p* < 0.05, Figure 5c). The limited macrophage recruitment observed following Omicron infection is consistent with its poor tropism for the OE and contrasts with the robust and spatially extensive inflammatory response induced by the Alpha variant.

### SARS-CoV-2 infection attenuates odor-evoked activity-dependent gene expression in mOSNs

Our scRNA-seq analysis identified reduced expression of several activity-dependent genes in mOSNs, suggesting that neuronal responsiveness to odor stimulation might be impaired at the periphery following SARS-CoV-2 infection. To directly test this hypothesis, we exposed Alpha-infected and mock-inoculated mice to an odor mixture and quantified the induction of activity-dependent genes in the OE. The efficacy of the odor mixture was first validated by measuring the levels of phosphorylation of ribosomal protein S6 (pS6), an established marker of odor-evoked OSN activation, in wildtype animals^43,44^ (supplemental Figure 5). However, because pS6 is regulated through the mTOR signaling pathway and can be influenced by cellular stress and antiviral signaling, it was not used as a quantitative readout in SARS-CoV-2 Alpha-inoculated animals. Instead, RNAscope in situ hybridization and immunohistochemistry were used to quantify the activity-dependent transcripts *S100a5*, *Kirrel2,* and LRRC3B protein^39,40^ in mock-inoculated and SARS-CoV-2 Alpha-inoculated mice with or without odor stimulation (Figure 6, n = 3 per group).

**Figure 6.**
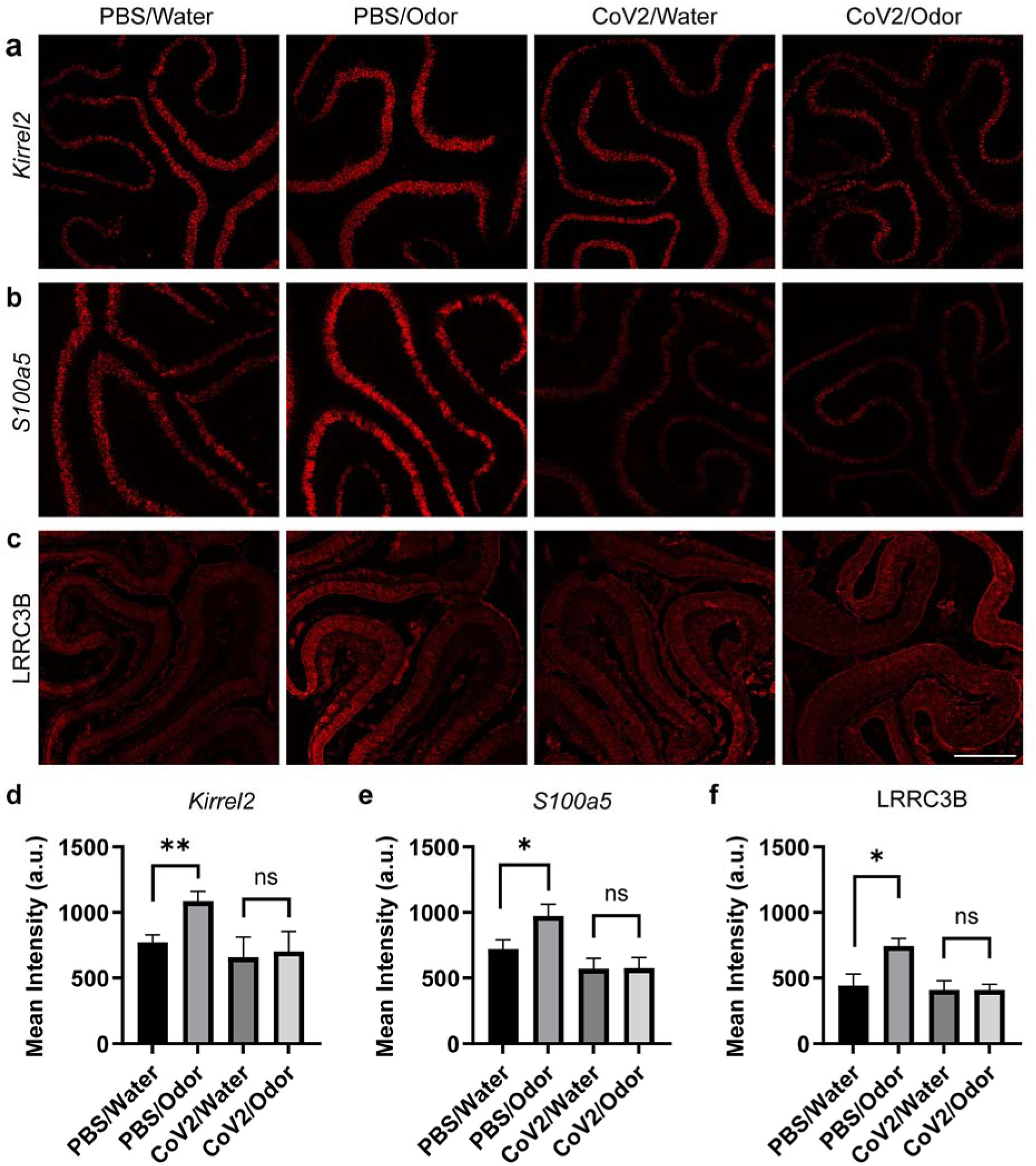
Odor stimulation induces mOSN activity-dependent gene expression in control but not infected mice. **a–c** RNAscope for *Kirrel2* (a) and *S100a5* (b) mRNA, and immunohistochemistry for LRRC3B (c) protein (red), in the OE of PBS mock-infected or Alpha-infected K18-hACE2 mice at 2 dpi, exposed to water (unstimulated) or odor mixture stimulation. Scale bars, 200 µm. **d–f** Quantification of mean staining intensity for *Kirrel2* (d), *S100a5* (e) and LRRC3B (f) in PBS/Water, PBS/Odor, CoV2/Water and CoV2/Odor groups. Odor stimulation increased marker expression in PBS mock-infected mice but not in Alpha-infected mice. Data are mean ± SEM; \**p* < 0.05, \*\**p* < 0.01, ns, not significant (Welch unpaired t-test).

Odor stimulation robustly induced transcript density in both activity-dependent genes, *Kirrel2* and *S100a5* in mock-infected animals from 772.9 ± 56.2 to 1086 ± 74.5 (Mean Intensity ± SEM, *p* < 0.006) and from 720.8 ± 71.4 to 974.4 ± 87.3 (*p* < 0.045) (Figure 6a,b,d,e). In striking contrast, odor stimulation failed to induce an increase in either transcript in SARS-CoV-2-inoculated animals. *Kirrel2* expression following water exposure was 658.8 ± 154.7, and following odor exposure was 701.6 ± 153.9 (*p* < 0.85). Similarly, *S100a5* expression following water exposure was 571.4 ± 77.08 and following odor exposure was 574.3 ± 80.90 (*p* < 0.98). Under water exposure conditions, expression of activity-dependent genes did not change significantly between SARS-CoV-2-infected animals and mock animals (*Kirrel2 p* < 0.51 and *S100a5 p* < 0.19).

Protein-level analysis of LRRC3B expression yielded similar results. Immunostaining of LRRC3B showed a significant increase upon odor stimulation in controls (441.0 ± 88.9 vs 743.9 ± 59.1, *p* < 0.02) (Figure 6c,f). Consistent with the transcript analysis, odor stimulation did not significantly upregulate LRRC3B protein expression in infected animals (*p* = 1.00). Together, these findings demonstrate that SARS-CoV-2 infection markedly impaired odor-evoked activity-dependent gene induction in mOSNs, indicating functional deficits in neuronal activation despite the absence of widespread neuronal loss.

## Discussion

In this study, we demonstrate that localized SARS-CoV-2 infection triggered a tissue-wide antiviral response throughout the OE despite no direct infection of OSNs. This widespread response induced a transient interferon-responsive state in mOSNs, followed by persistent suppression of mitochondrial, odor transduction, and activity-dependent gene programs. These transcriptional changes were accompanied by impaired odor-evoked neuronal activation despite preservation of the OSN population. Finally, comparison of SARS-CoV-2 variants, Alpha vs Omicron, demonstrated that antiviral activation and inflammatory remodeling correlated with local olfactory epithelial infection rather than systemic viral infection. Together, these findings identify tissue-wide bystander antiviral signaling as a mechanism linking localized SARS-CoV-2 infection to olfactory dysfunction.

While previous studies established that sustentacular cells are the principal targets of SARS-CoV-2 infection and proposed non-cell-autonomous mechanisms of olfactory dysfunction^18,19,23,45–47^, our findings directly visualize how localized infection is translated into tissue-wide antiviral signaling. Viral transcripts remained confined to focal infection sites, whereas ISG expression extended across neighboring sustentacular cells and OSNs in a spatial gradient, indicating widespread bystander interferon signaling. This response was largely absent following Omicron infection despite detectable pulmonary infection^31^. Omicron has been shown to have a reduced impact on anosmia in COVID-19 patients^48^. Though a hamster model showed that Omicron infected the OE and impacted olfaction^49^, we observed minimal SARS-CoV-2 infection in the OE of K18-hACE2 mice. This discrepancy may reflect species-specific differences in viral tropism in the OE between the animal models used. The difference in ISGs in the OE between Alpha- and Omicron-infected animals demonstrates that local OE infection, rather than systemic respiratory infection, was the principal driver of antiviral activation in the OE. These findings are consistent with the reduced incidence of olfactory dysfunction reported for Omicron infection and suggest that viral access to the olfactory mucosa is a key determinant of anosmia. Unlike the airway epithelium, the OE is a specialized neuroepithelial barrier that must simultaneously mount an antiviral defense while preserving neuronal function. The recently described blood–olfactory barrier may further reinforce the importance of local immune regulation by limiting access of circulating immune mediators to the olfactory mucosa^37,50^. Together, our findings support a model in which focal epithelial infection initiates a local antiviral program that propagates across the OE to reprogram neuronal function without widespread neuronal infection.

Our findings differ from reports describing widespread suppression of olfactory receptor genes following SARS-CoV-2 infection in hamsters and human tissue^51,52^. Instead, we found that olfactory receptor expression was largely preserved during early infection, whereas genes involved in odor signal transduction, mitochondrial metabolism, and activity-dependent plasticity showed the most prominent changes at 6 dpi^46,53^. This delayed or persistent response can also be seen with macrophage recruitment to the OE as a whole. While immune cell infiltration is an important component of the response to infection, its persistence and widespread increase raise the possibility that sustained immune activity contribute to OSN dysfunction despite minimal or resolving viral presence^54^. Consistent with the nuclear architecture model proposed previously, our findings suggest that interferon-mediated transcriptional reprogramming may represent an additional mechanism that contributes to olfactory dysfunction^46^.

Previous studies primarily attributed COVID-19-associated anosmia to epithelial injury and altered olfactory receptor expression. In contrast, our data demonstrate that mOSNs exhibited persistent suppression of activity-dependent gene programs and failed to mount transcriptional responses following odor stimulation. Phosphorylated ribosomal protein S6 (pS6) has been shown to be an excellent indicator of OSN activity. However, SARS-CoV-2 infection induces mTOR signaling, and thus pS6 could not be used to quantify OSN activity in this study. *S100a5*, *Kirrel2*, and LRRC3B are established activity-dependent genes whose expression reflects sensory-driven neuronal activation rather than neuronal survival^39,40,44,55^. Their impaired induction provides direct molecular evidence that surviving OSNs became functionally compromised after infection. These findings indicate that olfactory dysfunction can arise from impaired neuronal responsiveness in the absence of widespread neuronal loss and establish activity-dependent gene expression as a sensitive molecular readout of peripheral olfactory function during viral infection. Compared with pS6, which is regulated through the mTOR pathway and may be influenced by antiviral signaling and animal olfactory behavior test which does not pinpoint the origin of pathology, activity-dependent transcriptional responses provide a cell-type specific approach for assessing OSN function under inflammatory conditions.

Although heterogeneous hACE2 expression in the K18-hACE2 OE (supplement Figure 1) results in localized SARS-CoV-2 infection, this feature provides a useful platform for dissecting host responses to localized viral invasion. Future studies using this system should help identify the epithelial and immune signaling pathways that coordinate antiviral defense while preserving neuronal survival and determine whether these responses are primarily protective or contribute directly to neuronal dysfunction. Extending these analyses to human tissue and models of persistent olfactory dysfunction will further clarify how acute antiviral responses influence long-term recovery and may reveal therapeutic strategies to preserve or restore olfactory function following viral infection.

## Materials and Methods

### Mice

All mouse work was conducted based on the animal protocol (#23489) approved by the institutional animal care and use committee at the University of California, Davis. Infectious virus was handled in a certified animal biosafety level 3 laboratory (ABSL-3) spaces in compliance with approved institutional biological use authorization. All mouse work adhered to the NIH Guide for the Care and Use of Laboratory Animals.

Male and female K18-hACE2 (B6.Cg-Tg(K18-ACE2)2Prlmn/J) mice aged 7–12 weeks were purchased from The Jackson Laboratory (Jax # 034860). Mice were weighed and anesthetized with isoflurane, then inoculated intranasally via hanging drop over both nares with 30 μL DPBS or 10^4^–10^5^ PFU (total dose in both nares) of SARS-CoV-2 diluted in DPBS. Inocula were back-titrated by Vero plaque assay to confirm the dose. Mice were monitored for changes in weight and clinical signs of disease, including ruffled fur, ataxia, and labored breathing, twice daily until 6 dpi. Oropharyngeal swabs were collected under isoflurane anesthesia at 1 and 2 dpi. Mice were euthanized at prescribed time points on 1, 2, 4, and 6 dpi, followed by perfusion with sterile DPBS. The right inferior lobe of the lung and the left hemisphere of the brain or trachea were harvested for viral titer determination. Olfactory tissues were collected for experimental procedures.

### SARS-CoV-2 variants and titer determination

SARS-CoV-2 variant B.1 (human/USA/CZB-59 x 002/2020, GenBank #MT394528) isolated from a patient in 2020 in Northern California was provided by Dr. Christopher Miller. Alpha B.1.1.7 (hu/USA/CA_CDC_5574/2020, GISAID #EPI_ISL_751801), Beta B.1.351 (hCoV-19/USA/MD-HP01542/2021, GISAID #EPI_ISL_89-360), Delta B.1.617.2 (hCoV-19/USA/PHC658/2021, no GISAID number provided), and Omicron variants B.1.1.529 (hCoV-19/USA/HI-CDC-4359259-001/2021, GISAID: EPI_ISL_8690072) were obtained from Biodefense and Emerging Infections Research Resources Repository, National Institutes of Allergy and Infectious Diseases at the United States National Institutes of Health. All virus strains were propagated after procurement in Vero E6 or Vero CCL-81 cells to achieve titers >10^6^ PFU/mL and stored at -80 °C until use.

Infectious SARS-CoV-2 in tissue homogenates and oropharyngeal swab supernatants were determined by plaque assay. Serial dilutions of homogenates/supernatants were adsorbed onto confluent layers of Vero-E6 cells for 1 hour and then overlaid with agarose for 3 days. Plaques were visualized by crystal violet staining after fixing with 4% formaldehyde and viral titers were calculated as PFU per mg of tissue or per swab.

### Immunohistochemistry and in situ hybridization

Mice were transcardially perfused with PBS and subsequently fixed by immersion in 10% formalin at room temperature for 24–48 hours. Nasal tissues were decalcified in 0.45M EDTA in PBS for 3 days, cryoprotected in 30% sucrose, and embedded in OCT compound for cryosectioning. Immunohistochemistry was performed as previously described^31^. Primary antibodies used were: Rabbit anti-SARS-CoV-2 Nucleocapsid (Sino Biological, #40143-R01), Chicken anti-OMP (custom), Goat anti-Iba1 (Wako, #011-27991), Rabbit anti-LRRC3B (Novus, NBP1-89579). RNAscope in situ hybridization was performed according to the manufacturer’s protocol using Multiplex Fluorescent V2 Assay (Advanced Cell Diagnostics). Probes used are V-nCoV2019-S (#848561), Kirrl2 (#491421), S100a5 (#851971), Rsad2 (#561251), Isg15 (#559271), Ifit3 (#508251), Ifih1 (#1574241), Irf7 (#534541), Eif2ak2 (#822871).

### Bulk RNA-seq and qRT-PCR

Formalin-fixed olfactory mucosa from infected and control mice was dissected away from turbinates and homogenized using a TissueLyzer (Qiagen). Total RNA was extracted from triplicates of infected and control samples using a High Pure FFPE RNA Micro Kit (Roche, #04823125001). Following rRNA removal, RNA-seq libraries were prepared. Sequencing was performed on an Illumina NovaSeq 6000 platform. Raw sequencing reads were trimmed and filtered to remove low-quality reads and aligned to the reference mouse genome (GRCm38-mm10) using STAR v2.7. Differentially expressed genes were identified from the RNA-seq data using limma-voom with an adjusted p-value cutoff of 0.05 and a log2 fold-change threshold of ±1. GO enrichment analysis was performed using the enrichR package v3.4 with the org.Mm.eg.db annotation database. Gene-set analysis for selected GO terms was done using the QuSAGE method^38^.

For qRT-PCR, biological triplicates from PBS controls and infected mice at 1, 2, and 6 dpi were included. cDNA was synthesized from the total RNA using random hexamers and SuperScript IV (Invitrogen, #18090050). The initial cDNA was pre-amplified with a primer pool consisting of selected antiviral genes. qRT-PCR was performed using SybrGreen chemistry including technical triplicates. The differential gene expression was determined using the ΔΔCt method.

### scRNA-Seq and data analysis

Olfactory mucosae after fixation were dissected as described in the bulk RNA-seq procedure. Single cell suspension was obtained by mincing and digesting at 37°C with LiberaseTL (Roche) on a gentleMACS tissue dissociator (Miltenyi Biotec). After pelleting the dissociated cells at 900g for 5min, the supernatant was replaced with 2mL Quenching buffer (10x Genomics Next GEM Single Cell Fixed RNA Sample Preparation Kit, PN# 1000414) in DEPC-ddH2O to stop the Liberase reaction. The cell suspension was then filtered through a 40 um strainers and added 7-AAD dyes (Invitrogen, #A1310) for nuclei staining. Fluorescence-activated cell sorting was performed to obtain debris free single cell solution. Six samples of infected and control olfactory mucosae were loaded into Chromium Next GEM chips. Single cell RNA sequencing libraries were prepared using the Chromium Single Cell Gene Expression Flex v1 chemistry and sequencing was performed on Illumina NovaSeq platforms.

The raw single-cell Flex Gene Expression sequencing data was pre-processed using Cell Ranger multi pipeline v7.0 provided by 10X Genomics. The filtered gene expression data was imported to Seurat (v5) for further quality control. Cells were required to have the number of genes expressed in between 200 and 6,000, the number of unique transcripts expressed in between 500 and 40,000. Cell doublets were removed using scDblFinder.

Gene expression data were normalized using LogNormalize method in Seurat with a scale factor of 10000. PCA technique was used for dimensionality reduction to 50 PCs. Batch correction was performed with Harmony using group.by.vars = “sample”, theta = 2, lambda = 1, correcting simultaneously for per-individual mouse variation and sequencing batch effects. Clustering and UMAP were generated using the top 25 PCs. Cell type identification was carried out on clusters generated at resolution 0.4, using curated marker genes. Pseudobulk differential expression (DE) analysis was done using DESeq2. Significance was defined as BH-adjusted *p* < 0.05.

Pathway enrichment analysis was performed on significantly regulated genes per cell type from the DE results. Pathway enrichment analysis was performed using enrichGO (clusterProfiler, org.Mm.eg.db) with BH correction. Semantic redundancy was reduced using simplify() applied per cell type (Wang method, cutoff = 0.7) on each enrichR object individually before combining across cell types.

mOSNs identified in annotated object were subsetted and processed independently of the other cell types. Resolution 0.4 was selected as the working subcluster resolution. All mOSN cells were scored on a 24-gene canonical mouse interferon-stimulated gene panel (*Irf7, Isg15, Ifit1/2/3, Oasl1/2, Oas1a/2/3, Rsad2, Mx1/2, Stat1/2, Ifi27, Usp18, Cxcl10, Ifih1, Ddx58, Bst2, Irf9, Zbp1, Isg20*) using Seurat::AddModuleScore(). ISG-high threshold was calibrated against the SARS-CoV-2-dominant clusters’ own score distribution. Two tiers were defined, ISG-high and ISG-low, for mOSNs of the infected samples. DE (Seurat::FindMarkers, Wilcoxon rank-sum test on single cell expression, Bonferroni-adjusted) and gene set enrichment analysis were performed for each tier and activity dependent genes.

To enable a real cross-timepoint comparison, a raw count from both timepoints’ mOSN object were merged into one object and normalized. AddModuleScore was run across the pooled cells from both timepoints. This gives one shared control-gene background, so scores are genuinely comparable across timepoints within a panel. Two metrics were computed per panel and timepoint: breadth – the percentage of CoV2 cells scoring above threshold and amplitude – the median score among only those above-threshold (induced) cells.

### Odorant-induced olfactory sensory neuron activation

The odor mixture used to activate olfactory sensory neurons contained acetophenone, β-citronellol, muscone, and methyl tigilate in equal proportions. Twenty-four hours before odor exposure, mice were transferred to cages containing Alpha-Dri bedding and housed individually. One hour before exposure, mice were transferred to new cages to ensure a clean odor environment. Odorant mixture (10 uL) was placed on a 1 x 1 cm2 filter paper. The filter paper was then secured in a perforated cassette and placed inside the cage. Animals were exposed to the odorant mixture for 1 hour, after which they were euthanized, and tissues were collected.

Activity-dependent gene expression, together with *Omp* and either *Isg15* or *Rsad2*, was visualized by RNAscope in situ hybridization, and images were acquired by confocal microscopy. Signal quantification for activity-dependent genes was performed using ImageJ. All experimental groups contained biological triplicates. For mock control OEs, regions of interest (ROIs) were defined by automatically thresholding the *Omp* signal in ImageJ. The integrated density of the activity-dependent gene signal within each ROI was measured and normalized to the ROI area. For SARS-CoV-2-infected samples, antiviral response-positive, *Omp*-positive regions were thresholded in ImageJ and used to define ROIs. The integrated density of the activity-dependent gene signal within these ROIs was then calculated and normalized by the ROI area, as described for the PBS control condition. Raw data were imported into GraphPad Prism for statistical analysis, where unpaired Welch t-tests were performed to compare groups.

### Macrophage Infiltration

Macrophage presence (Iba1+) and infection site (NP+) were visualized by immunostaining, and images were acquired by confocal microscopy. Percent area was calculated using ImageJ. All experimental groups contained biological triplicates. ROIs were sampled from three specific areas: non-infected loops (no NP present), infected loops (NP present), and the septum. For mock control OEs, ROIs were taken from similarly localized areas where infection was seen in SARS-CoV-2-infected OEs. Within each ROI, the Iba1 signal area was normalized against the total ROI area to get the percentage of Iba1. Technical triplicates from each sampled area were averaged and then compared across biological triplicates for statistical analysis. Raw data were imported into GraphPad Prism for statistical analysis, where unpaired t-tests were performed to compare selected areas to their respective non-infected controls.

## Data availability

The bulk RNA-sequencing data generated in this study (whole olfactory epithelium, 24 and 48 h post-infection) have been deposited in the Gene Expression Omnibus (GEO) under accession number GSE345081. The single-cell RNA-sequencing data generated in this study (olfactory epithelium, 1 and 6 days post-infection) have been deposited in GEO under accession number GSE347396. Source data are provided with this paper.

## Acknowledgement

We thank the UC Davis DNAtech core for their expert support in transcriptomic studies, Dr. Richard Tucker, and all lab members for their constructive criticism and critical reading of the manuscript. We thank Dr. Paul Feinstein for his expert suggestions.

## Author contributions

QG and LLC designed the study. JL, MSA, HL, BMR, AC, and MC performed the experiments. ALW, JL, BMR, HL, and YK analyzed the data. JL, QG, and MSA wrote the manuscript. LLC and SR provided critical input and revised the manuscript. All authors reviewed and approved the final manuscript.

## Supplemental Material

**supplemental Figure 1.**
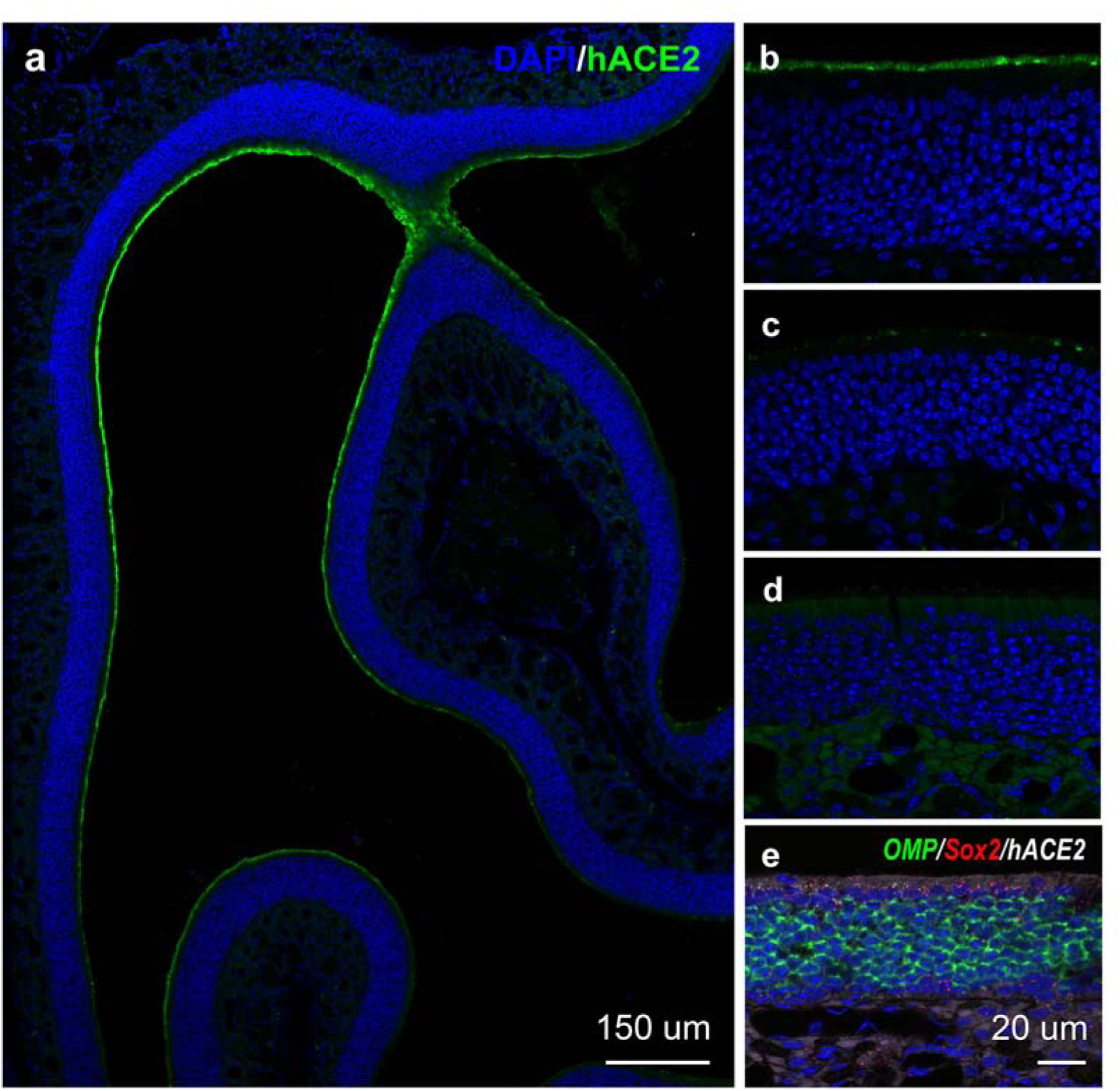
Human ACE2 is expressed by sustentacular cells in the K18-hACE2 mouse olfactory epithelium. **a** Low-magnification image of a coronal section through the nasal turbinates of a K18-hACE2 mouse showing hACE2 immunostaining (green) concentrated along the apical surface of the epithelium, with heterogeneous signal intensity across turbinate loops; nuclei counterstained with DAPI (blue). **b–c** Higher-magnification images of representative turbinate regions illustrating the variable intensity of hACE2 immunosignal (green) across different areas of the OE; nuclei (DAPI, blue). **d** hACE immunosignal is lacking in wildtype C57BL/6J OE. **e** High-magnification multiplex RNAscope image showing co-expression of hACE2 (white) with Sox2 (red), a sustentacular cell marker, together with OMP (green) marking olfactory sensory neurons and DAPI (blue) counterstaining nuclei, demonstrating that hACE2 is expressed by sustentacular cells rather than OSNs. Scale bars, 150 µm (a) and 20 µm (b-e).

**supplemental Figure 2.**
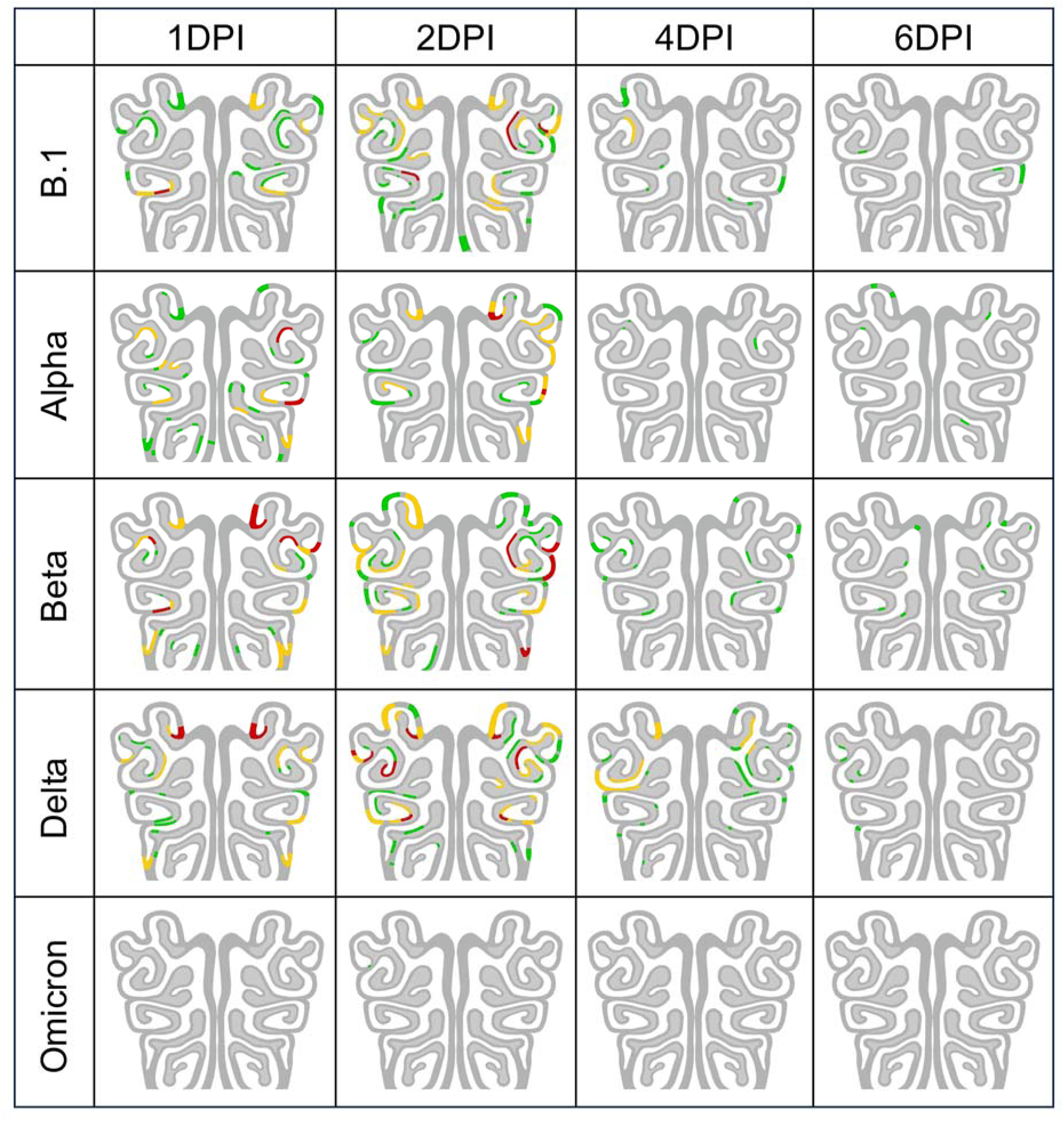
Spatiotemporal resolution of SARS-CoV-2 infection in the olfactory epithelium across variants and time. Heatmaps of nasal epithelium infection scores for each SARS-CoV-2 variant (rows: B.1, Alpha, Beta, Delta, Omicron) at 1, 2, 4 and 6 dpi (columns; n = 3 mice per variant per timepoint). Average NP infection score per region of interest is color-coded as in Figure 1h: green, score =1; yellow, score = 2; red, score = 3. Multi-cell infected clusters were largely resolved by 4 dpi and fully resolved by 6 dpi for all variants tested; Omicron showed no detectable OE infection at any timepoint.

**supplemental Figure 3.**
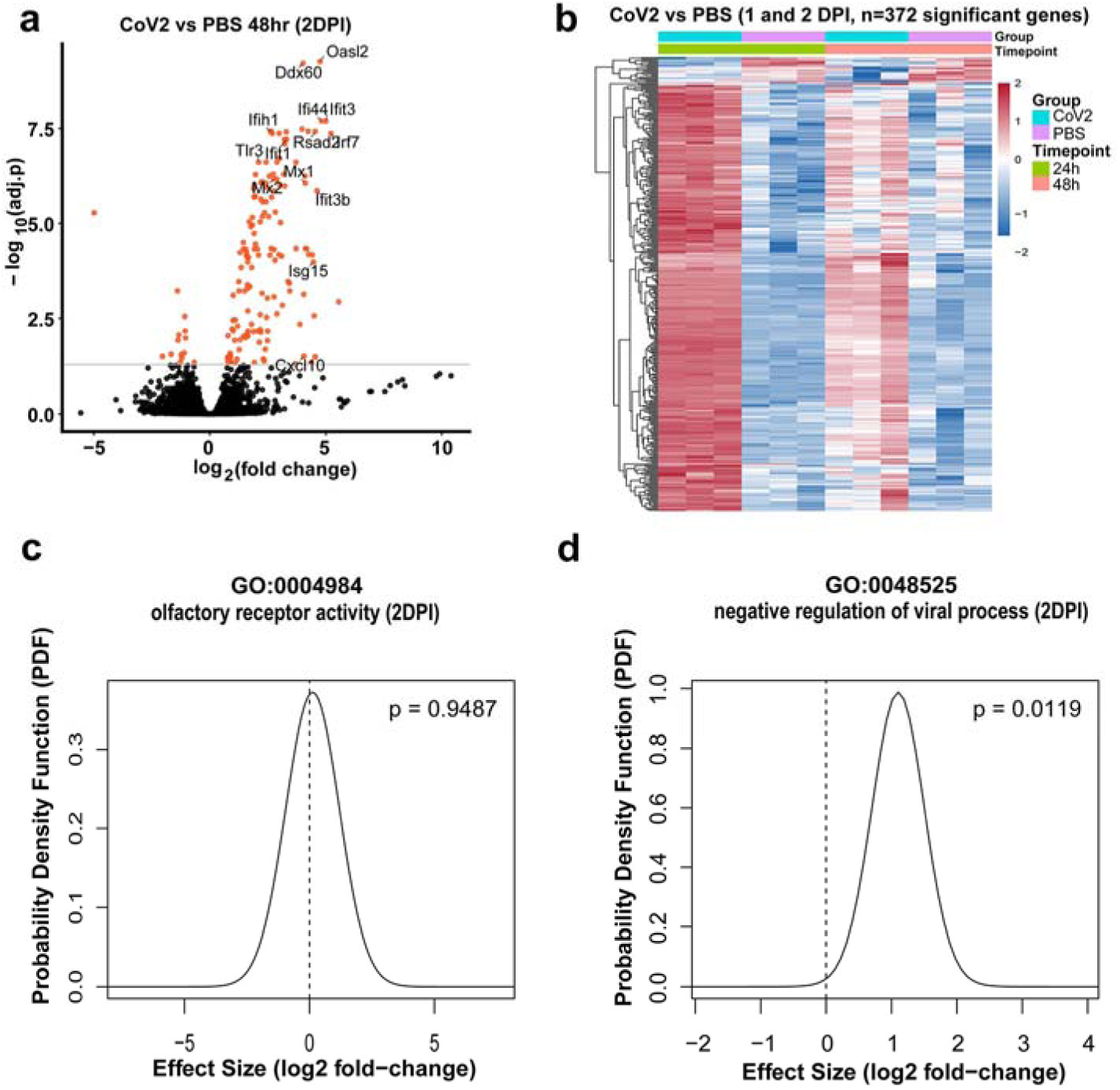
Transcriptional response to SARS-CoV-2 infection is attenuated by 2 dpi. **a** Volcano plot of differentially expressed genes in bulk olfactory mucosa from SARS-CoV-2 (Alpha)-infected versus PBS mock-infected K18-hACE2 mice at 2 dpi (n = 3 per group); selected interferon-stimulated genes are labeled. **b** Heatmap of relative expression (row z-score) of differentially expressed genes at 1 and 2 dpi, SARS-CoV-2 versus PBS (n = 3 per group per timepoint). **c, d** QuSAGE gene-set analysis of GO:0048525, negative regulation of viral process (c), and GO:0004984, olfactory receptor activity (d), at 2 dpi. Curves show the probability density function (PDF) of the gene-set effect size (log2 fold-change) relative to the null (dashed line); olfactory receptor activity gene set showed no significant shift at 2 dpi (*p* > 0.5), and negative regulation of viral process gene set showed a significant but reduced upregulation compared to 1 dpi (Figure 2d).

**supplemental Figure 4.**
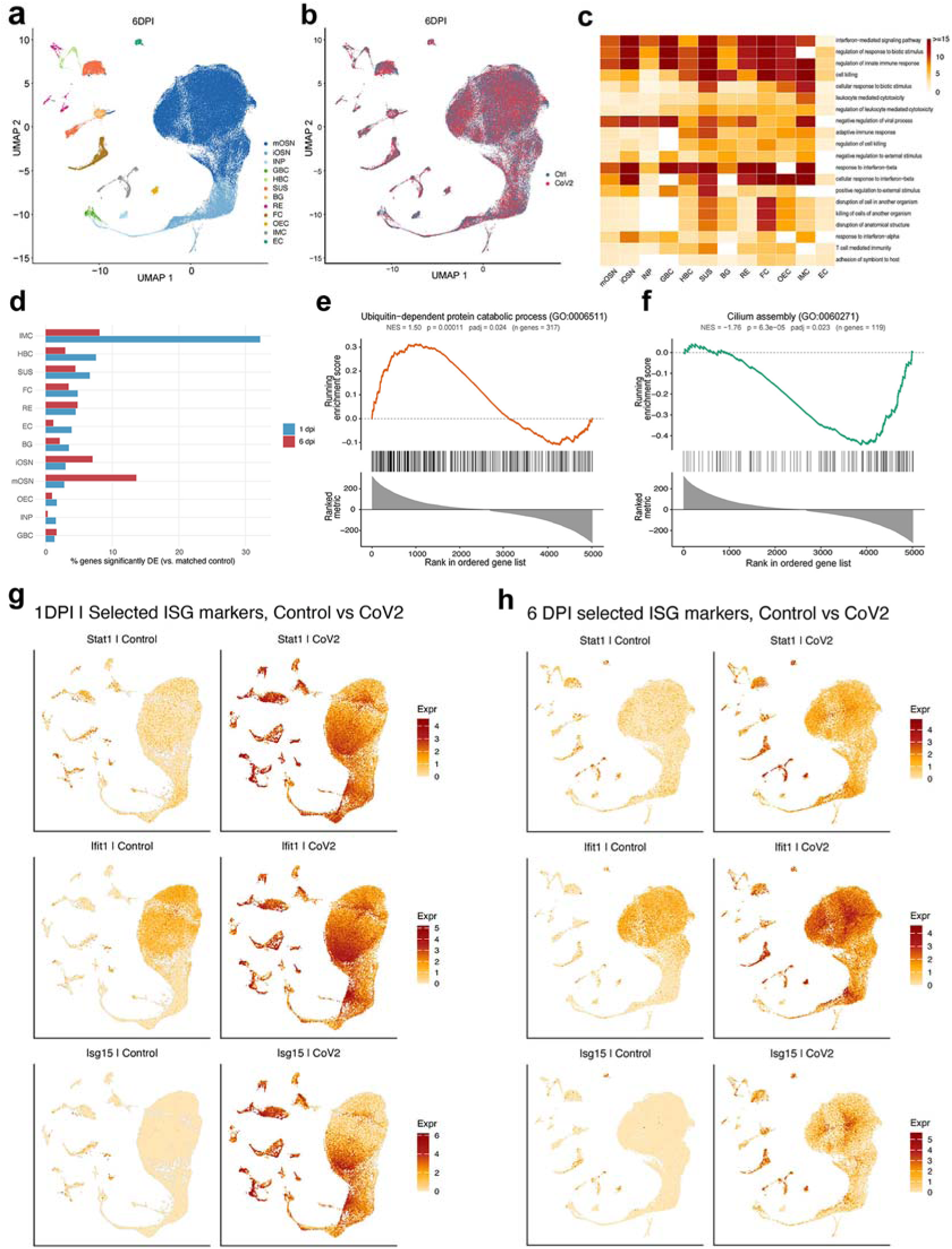
Response to SARS-CoV-2 infection persists to 6 dpi. **a** UMAP of olfactory mucosa scRNA-seq at 6 dpi, colored by cell type (as in Figure 4a). **b** UMAP colored by condition (control, CoV2). **c** Heatmap of Gene Ontology biological process terms enriched among upregulated differentially expressed genes per cell type, simplified per cell type (Wang similarity 0.7) and colored by −log10(adj.*p*), complementing Figure 4c. **d** Percentage of genes significantly differentially expressed relative to matched controls, by cell type, at 1 dpi (blue) and 6 dpi (red). **e, f** Gene set enrichment analysis (GSEA) at 6 dpi for ubiquitin-dependent protein catabolic process (GO:0006511) (e) and cilium assembly (GO:0060271) (f). Running enrichment score (top), gene hits along the ranked list (middle) and the ranked-list metric (bottom) are shown; normalized enrichment score (NES), nominal P and BH-adjusted P are indicated. **g, h** UMAP feature plots of representative interferon-stimulated genes (*Stat1, Ifit1, Isg15*) in control and SARS-CoV-2-infected olfactory mucosa at 1 dpi (g) and 6 dpi (h); color scale indicates log-normalized expression.

**supplemental Figure 5.**
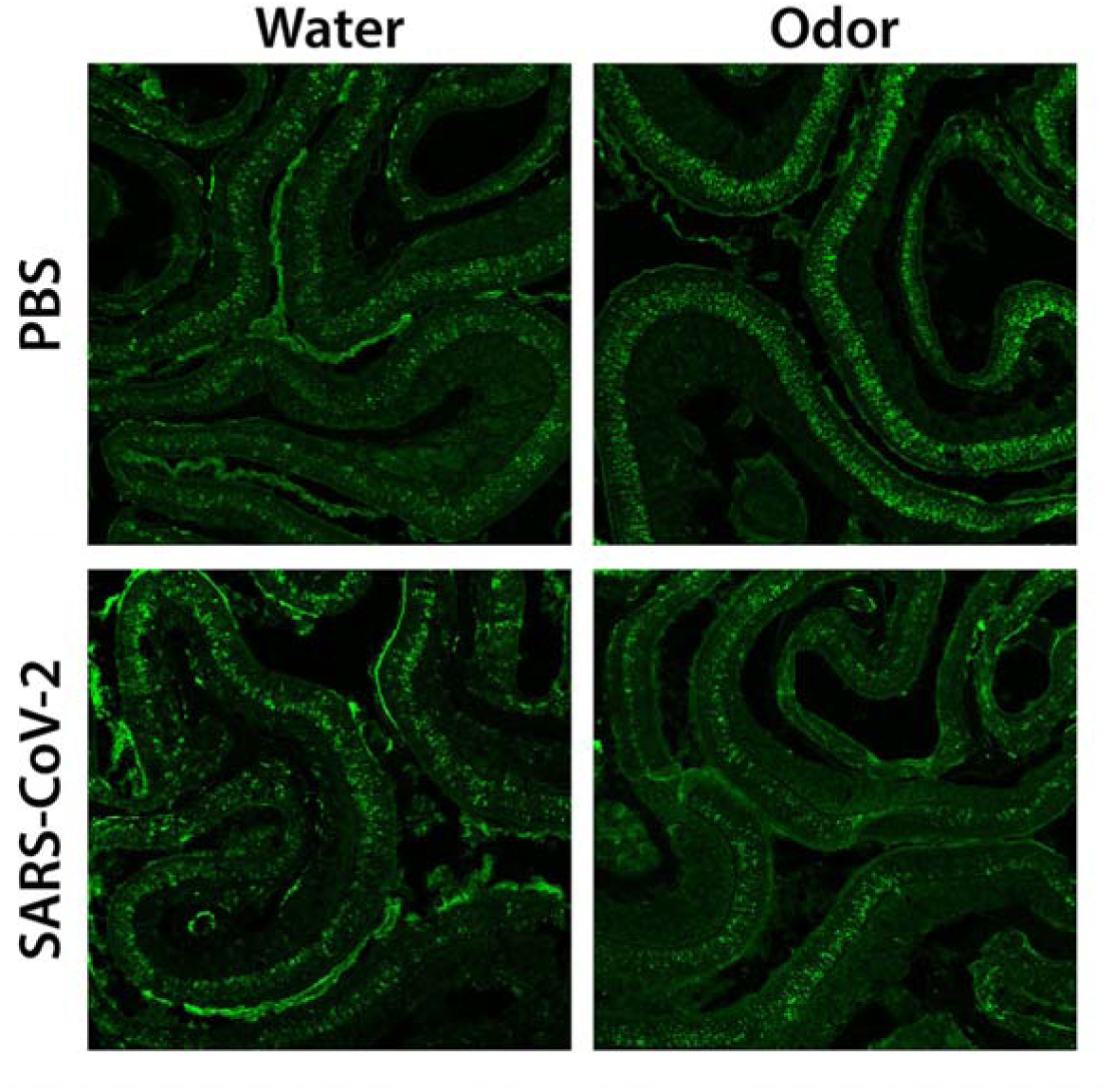
Odor stimulation induces phospho-S6 (pS6) signal in the olfactory epithelium. Representative immunofluorescence images of pS6 (green) in the OE of PBS mock-inoculated (top) and SARS-CoV-2 (Alpha)-infected (bottom) mice following water (control, left) or odor mixture exposure (right), confirming that the odor mixture activates OSNs.

**Supplemental Figure 6.**
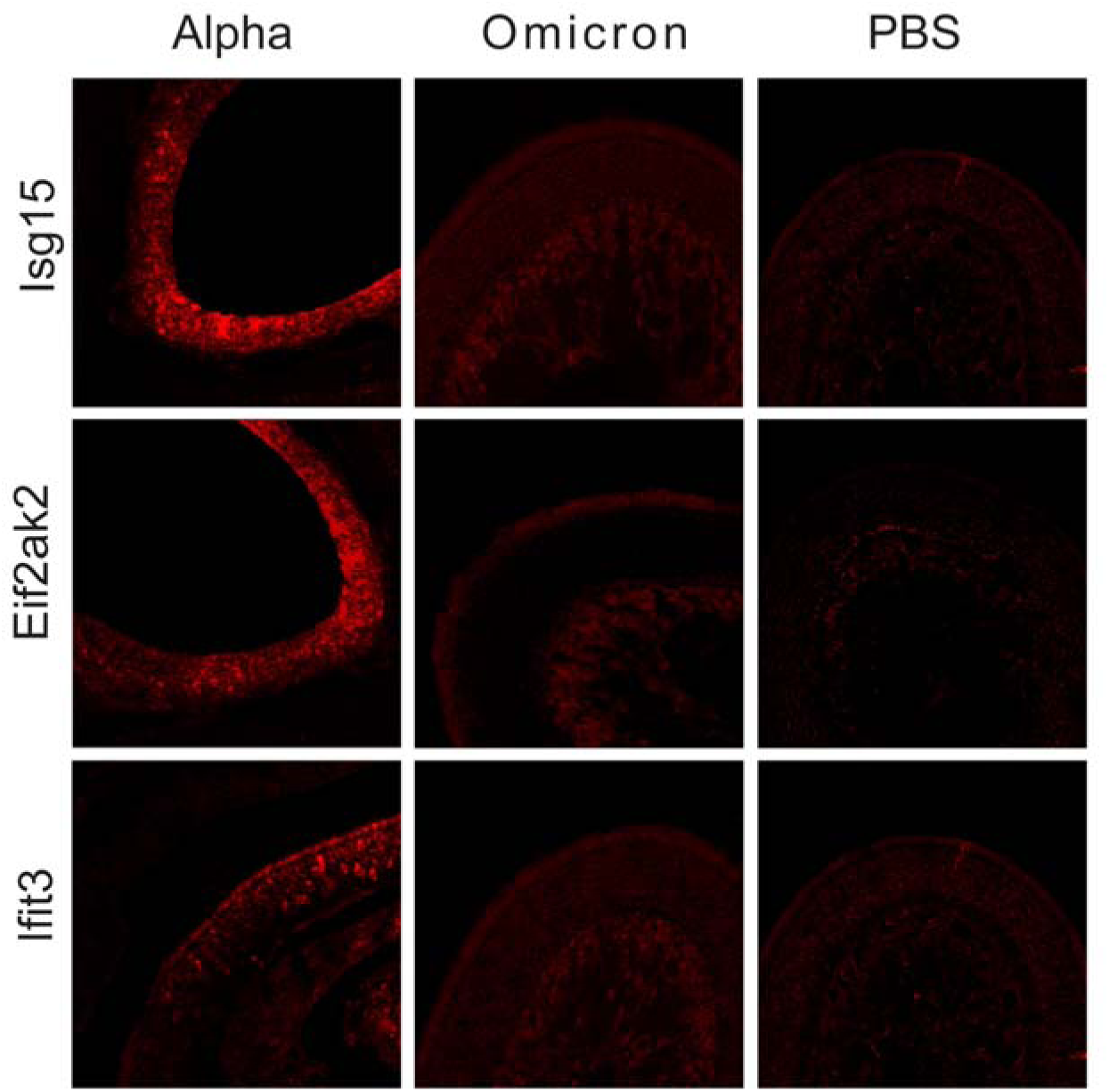
Comparison of antiviral gene expression, including *Isg15*, *Eif2ak2*, *Ifit3*, among OE tissues infected with the SARS-CoV-2 Alpha or Omicron variant or PBS mock control.

